# Sensitivity in the antioxidant system of discus fish (*Symphysodon* spp.) to cold temperature: evidence for species-specific cold resistance

**DOI:** 10.1101/749705

**Authors:** Shi-Rong Jin, Bin Wen, Zai-Zhong Chen, Jian-Zhong Gao, Lei Wang, Ying Liu, Han-Peng Liu

## Abstract

The discus fish (*Symphysodon* spp.) is an endemic species of the Amazon that is among the most popular ornamental fish around the world, and is usually used as the model animal for studying the diversification of Amazon fish. Here, a comparative analysis of two species of discus fish, i.e., *S. haraldi* and *S. aequifasciatus*, based on several antioxidant indexes was conducted, to test the hypothesis that cold resistance might correlate with the diversification of discus fish. We set up a continuous sequence of three temperature programs, namely cooling (28 °C to 14 °C; -1 °C/h), cold maintenance (14 °C for 12 h) and recovery (14 °C to 28 °C; +1 °C/h). Subordinate function (SF) combined with principal component analysis (PCA) showed that the cold hardiness of *S. haraldi* during cold treatment was in the order of cooling > cold maintenance ≈ recovery, but the cold hardiness of *S. aequifasciatus* during cold treatment was in the order of cold maintenance > cooling > recovery. Specifically, the lowest cold hardiness was observed in *S. aequifasciatus* during recovery, indicating that cold stress resulted in more seriously oxidative stress in *S. aequifasciatus* than in *S. haraldi*. Overall, these results show a significant interspecific variation, indicating the correlation between environmental adaptation and the diversification of discus fish.

## 1. Introduction

The discus fish (*Symphysodon* spp.) is an important ornamental tropical fish all over the world, originating from the Amazon River (Wen et al., 2018, 2017). In addition to their Amazon basin-wide distribution, the 3 currently recognized species of the genus *Symphysodon* (*S. aequifasciatus*, *S. discus*, and *S. haraldi*, Cichlidae, Perciformes) (Bleher et al., 2007; Gross et al., 2009) exhibit a large amount of morphological variation (different color and color patter) and genetic variability associated with different types of biotopes (Farias and Hrbek, 2008; Koh et al., 1999). For example, *S*. *haraldi*, the ‘bule’ discus, is found in the central portion of the Amazon basin (type locality Manacapuru river), *S. aequifasciatus*, the ‘green’ discus, is found in the west portion of the Amazon basin (type locality Tefé River), and *S. discus*, the. Heckel discus, is found in the Negro River basin (Farias and Hrbek, 2008; Gross et al., 2010).

Fish as an ectotherm, ambient temperature which constrains whole-organism performance is one of the most important factors affecting the biogeographic distribution and abundance (Troia and Gido, 2017). Recently, several studies have shown that distinct responses of antioxidant defense systems (ADS) would occur between different locations of fish species toward temperature stress (e.g., Bryant et al., 2018; Chung et al., 2017; Johnston et al., 1998; Rudneva-Titova et al., 1994 Shaliutinakolešová et al., 2013). Yet surprisingly few studies have compared thermal performance among closely related warm-water species (Troia and Gido, 2015).

We have a hypothesis that, depending on the geographic region and due the evolution, different species of discus fish may have different sensitivity to cold stress. To examine whether species-specific cold hardiness of discus fish, we exposed two species of discus fish (*S. haraldi* and *S. aequifasciatus*) to an acute cold stress, including a rapid temperature decrease (from 28 to 14° C), maintained up to 12 h, and then rapidly increase (from 14 to 28° C). ROS generation, ADS together with oxidative damage production were measured with the assumption that they were indicators of cold hardiness. Comprehensive evaluation using subordinate function (SF) method combined with principal component analysis (PCA) revealed cold hardiness of different species of discus fish.

## 2. Materials and methods

### 2.1 Experimental design

Juvenile discus fish (*S. haraldi* and *S. aequifasciatus*, body weight 9.24±1.63 g) were obtained from the Ornamental Fish Breeding Laboratory, Shanghai Ocean University (Shanghai, China). Then, 60 juvenile fish of each species were randomly divided into 3 glass aquaria (150 L), and acclimated at a temperature of 28 °C for a period of 30 days before the temperature trial. After acclimation, all fishes were subjected to a continuous sequence of three thermal treatments (namely cooling, cold maintenance and recovery) each. A schematic representation of the experimental procedure is provided in Fig. 1. First, the temperature in six aquaria initially decreased from 28 °C to 14 °C by 1 °C/h (cooling) and 9 fish (3 fish/aquaria) of each species were randomly sampled at t1 (28 °C), t2 (21 °C) and t3 (14 °C), respectively. When the coldest experimental temperature of 14 °C was attained, then fish were maintained at 14 °C for 12 h (cold maintenance) and other 9 specimens (3 fish/aquaria) of each species were collected at t4 (6 h) and t5 (12 h), respectively. At the end of experiment period, the remaining fish returned from 14 °C to the initial temperature value of 28 °C by 1 °C/h (recovery) and all were finally sacrificed and sampled at t6 (5 fish/aquaria; 15 fish totals per species). All animal care was conducted in accordance with the Administrative Measures for Experimental Animals in Shanghai, and the experimental protocols were approved by the Animal Ethics Committee of Shanghai Ocean University (SHOU IACUC protocol # 20171015).

**Fig. 1.**
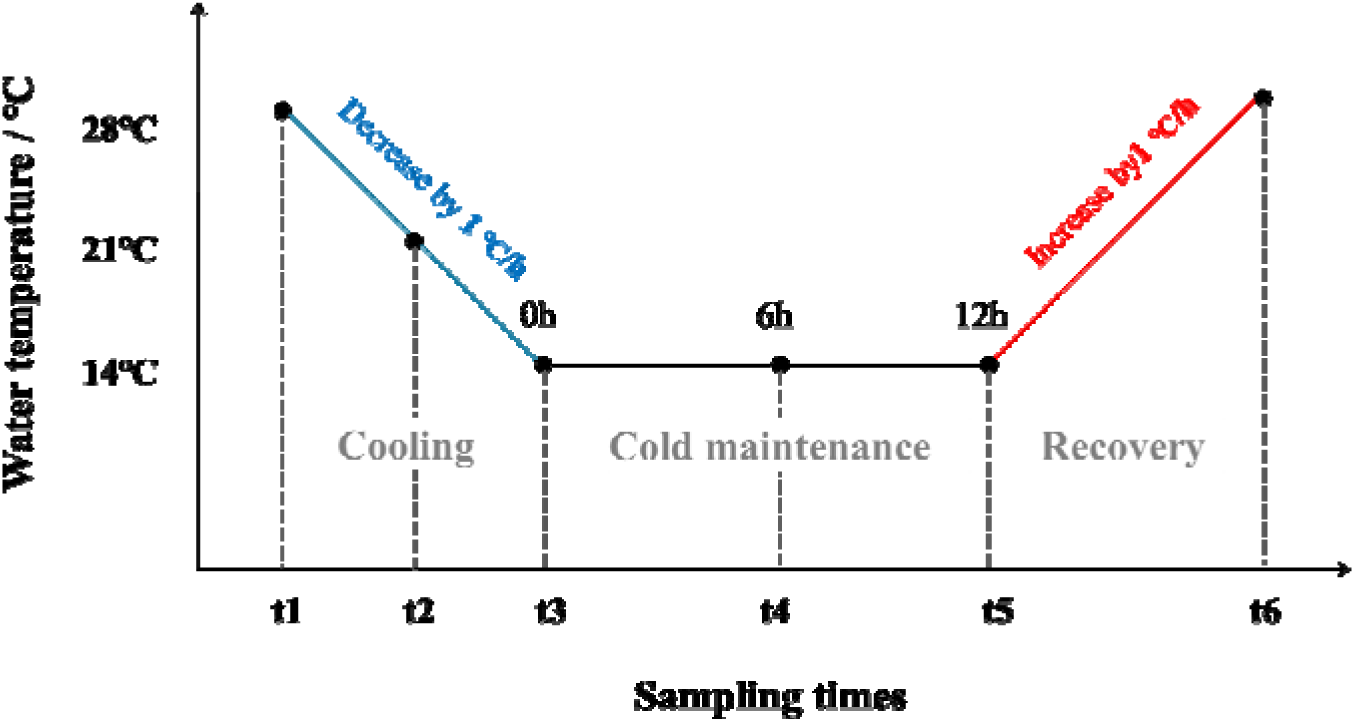
Scheme of the experimental design. Water temperature changes (solid line) and sampling times (t1-t6).

### 2.2 Sampling procedures

At each sampling times, the gill tissues were excised from each fish, rapidly deep-frozen in liquid nitrogen and stored at −80 °C for further analyses. On the one hand, gill samples were filtered by 300 mesh screens after homogenized (1:2, w/v) in ice-cold phosphate buffer (PBS; 0.1 M, pH 7.4). The single cell suspensions were used to measure the reactive oxygen species (ROS) content. On the other hand, the sampled gill tissues were homogenized (1:9, w/v) in an ice-cold NaCl 0.7% solution. The obtained homogenates were centrifuged at 3000 g at 4 °C for 10 min, and the supernatant were collected for other analysis.

Indicator tests were performed using classical colorimetric methods with commercial kits (Nanjing Jiancheng Bioengineering Institute, Nanjing, China). All assays, except ROS, were quantified spectrophotome trically, with a Synergy H4 Hybrid Multi-Mode microplate reader (BioTek Instruments, Winooski, VT, USA).

### 2.3 Reactive oxygen species (ROS) contents analysis

ROS level was measured using 2, 7-dichlorodihydrofluoresce in diacetate (H2DCFDA; Molecular Probes, Nanjing Jiancheng Bioengineering Institute, Nanjing, China), which was oxidized to fluorescent dichlorofluorescein (DCF) by intra-cellular ROS. A BD Accuri™ C6 flow cytometer (BD Biosciences) was used to analyze the ROS content. Data were recorded as cell cytograms showing the granularity (SSC value), relative size (FSC value), and fluorescent channels for each parameter. Each sample analysis included a total of 20,000 events, and the flow speed was maintained at less than 300 s^−1^. FL1 fluorescent channel were set to evaluate it (Wang et al., 2012).

### 2.4 Antioxidant enzymatic measurements

Total superoxide dismutase (SOD) activity was measured at 550 nm using the xanthine oxidase method that the protein amount giving 50% inhibition of maximum colour development contained 1 unit (U) of SOD (McCord and Fridovich, 1969). The results are accordingly given as U SOD/mg protein.

Catalase (CAT) activity was based on the reaction of the enzyme with methanol in the presence of an optimal concentration of H2O2 (Johansson and Borg, 1988). The purple color formed in these reactions was measured at 405 nm to measure CAT activity.

The activity of glutathione peroxidase (GPx) was measured at 412 nm, because GSSG occurs in the medium reduced to GSH by GPx and rate of GSH oxidation was used to calculate GPx activity (Hafeman et al., 1974). It calculated in terms of decreasing in GSH concentration by 1 μmol/L as one unit of enzyme activity.

Glutathione reductase (GR) activity was indirectly determined by mea-suring nicotinamide adenine dinucleotide phosphate (NADPH) consumption. Then the decrease in NADPH absorbance at 340nm was measured with a spectrophotometer (Carlberg and Mannervik, 1975).

Glutathione-S transferase (GST) activity was assayed by following the formation of glutathione–chlorodinitrobenzene (CDNB) adduct at 412 nm by the decreasing in reduced GSH concentration (Habig et al., 1974).

### 2.5 Glutathione contents analysis

The level of reduced Glutathione (GSH) was measured at 412 nm by using 5,5′-dithiobis(2-nitrobenzoic acid) (DTNB) reagent, following the method of Tietze (1969). DTNB was reduced by the free sulfhydryl groups of GSH to form the yellow compound 5-thio-2-nitrobenzoic acid (TNB).

### 2.6 Oxidative damage measurements

Malondialdehyde (MDA) and protein Carbonylation (PC) contents are relatively direct indexes for low-temperature damage (Ren et al., 2018; Vinagre et al., 2012; Ye et al., 2016). The higher content, the lower hardiness showed.

PC was measures via a reaction with 2,4-dinitrophenylhydrazine DNPH followed by TCA precipitation as previously described (Levine et al., 1994; Reznick and Packer, 1994).

MDA occurs in lipid peroxidation and this is measured after incubating at 95 °C with thibabituric acid (TBA) in aerobic condition (pH 3.4) (Uchiyama and Mihara, 1978). The pink colour formed in these reactions is measures in the spectrophotometer at 532 nm to measure MDA levels (Ohkawa et al., 1979).

### 2.7 Statistical analyses

For all parameters, data were expressed as mean ± standard error (SE). Statistical analyses were performed using SPSS, PASW statistics 20.0. A one-way analysis of variance (ANOVA) was performed to test for the effects of sampling species and temperature on the oxidative stress response values (tested separately) followed by the Tukey test. A significance level of 0.05 was used in all test procedures.

To intuitively inspect the tendencies in the variation of hardiness indexes between the temperature processing and species, we produced a heat map and comprehensively evaluated hardiness indexes using subordinate function (SF) method combined with principal component analysis (PCA) and correlation analysis.

Follow the data were standardized by Z-score, the heatmap was constructed by Sanger (V1.0.9).

Principal component analysis (PCA) and correlation analysis were performed using SPSS.

Subordinate function values were calculated, and average membership and cold resistance of different discus species were analyzed according to Zhang et al. (2015) and Zhao et al. (2019)

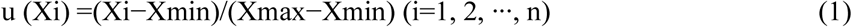

Weights of various comprehensive indicators were calculated as:

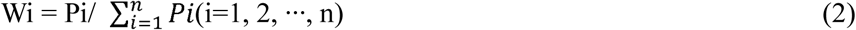

The D values of the different treatments were calculated as:

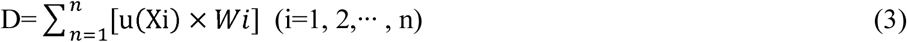

Xi, Xmin, Xmax and W□ are the score, minimum score, maximum scores and importance (weight) of the ith comprehensive indicator, respectively; Pi is the contribution rate of the □th comprehensive indicator of various treatments of two species; D is the comprehensive evaluation value for adaptability.

## 3. Results

### 3.1 Reactive oxygen species (ROS)

ROS level showed a slight fluctuation in *S. aequifasciatus* from t1- t5 (cooling and cold maintenance) until it reached the lowest value, then it showed a sharp rise during recovery. On the contrary, ROS level showed a sharp rise first (t1-t2) in *S. haraldi*, then progressively decrease and recovered to initial level. Due to high individual variability, the latter value was always statistically significantly higher than the former except t6 (*p* < 0.05) (Fig. 2).

**Fig. 2.**
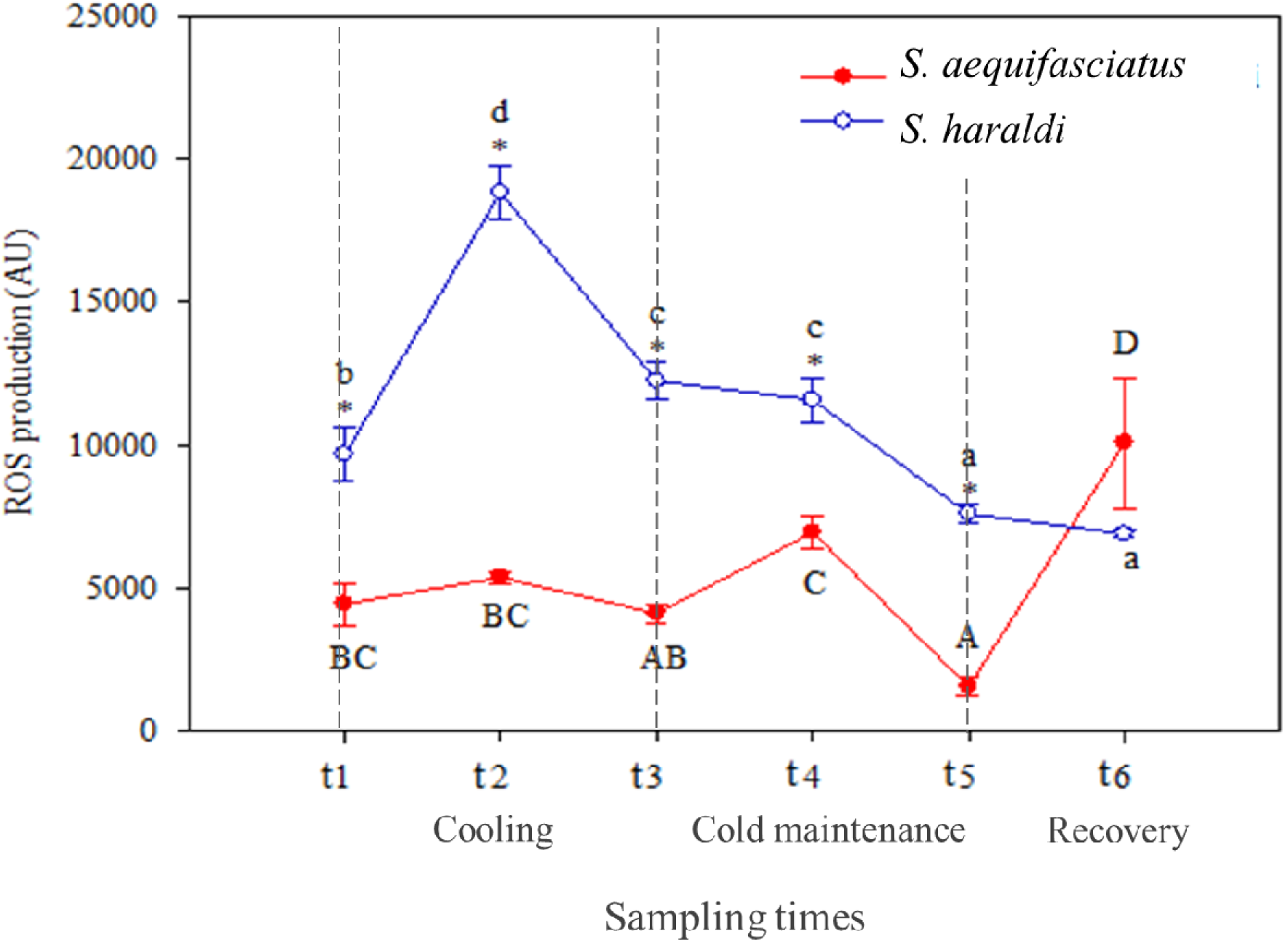
Production of reactive oxygen species in the gills of two subspecies of *Symphysodon spp*. red line, *S. aequifasciatus* and blue line, *S. haraldi*. Data are presented as means ±SD (n=3). *indicates significant differences (*p<*0.05) between subspecies. Different uppercase letters indicate significant differences (*p<*0.05) between sampling times within the *S. aequifasciatus*. Different lowercase letters indicate significant differences (*p<*0.05) between sampling times within the *S. haraldi*.

### 3.2 Antioxidant enzymatic activities

The temperature effects on antioxidant enzymatic system in two species are represented in Fig. 3. There were no effect of temperature on SOD (Fig. 3a) and GR (Fig. 3b) activity in *S. haraldi*, but in *S. aequifasciatus*, only CAT (Fig. 3c) activity was not affected. During cooling, SOD activity and GPx (Fig. 3d) activities showed an increase trend in *S. aequifasciatus*. In *S. haraldi*, CAT activity increased first and then recovered, while GPx activity decreased first and then significant increased during cooling. During cold maintenance, GPx and GR activities increased in *S. aequifasciatus*, but SOD activity showed decrease. Oppositely, there were no affects in *S. haraldi* during cold maintenance, except GPx activity showed fluctuation. During recovery, in both species, all antioxidant enzymatic activities except GST were recovered. GST activity was progressively increased in both species throughout the experiment (Fig. 3e), and finally *S. aequifasciatus* value was higher (*p* < 0.05) than *S. haraldi*.

**Fig. 3.**
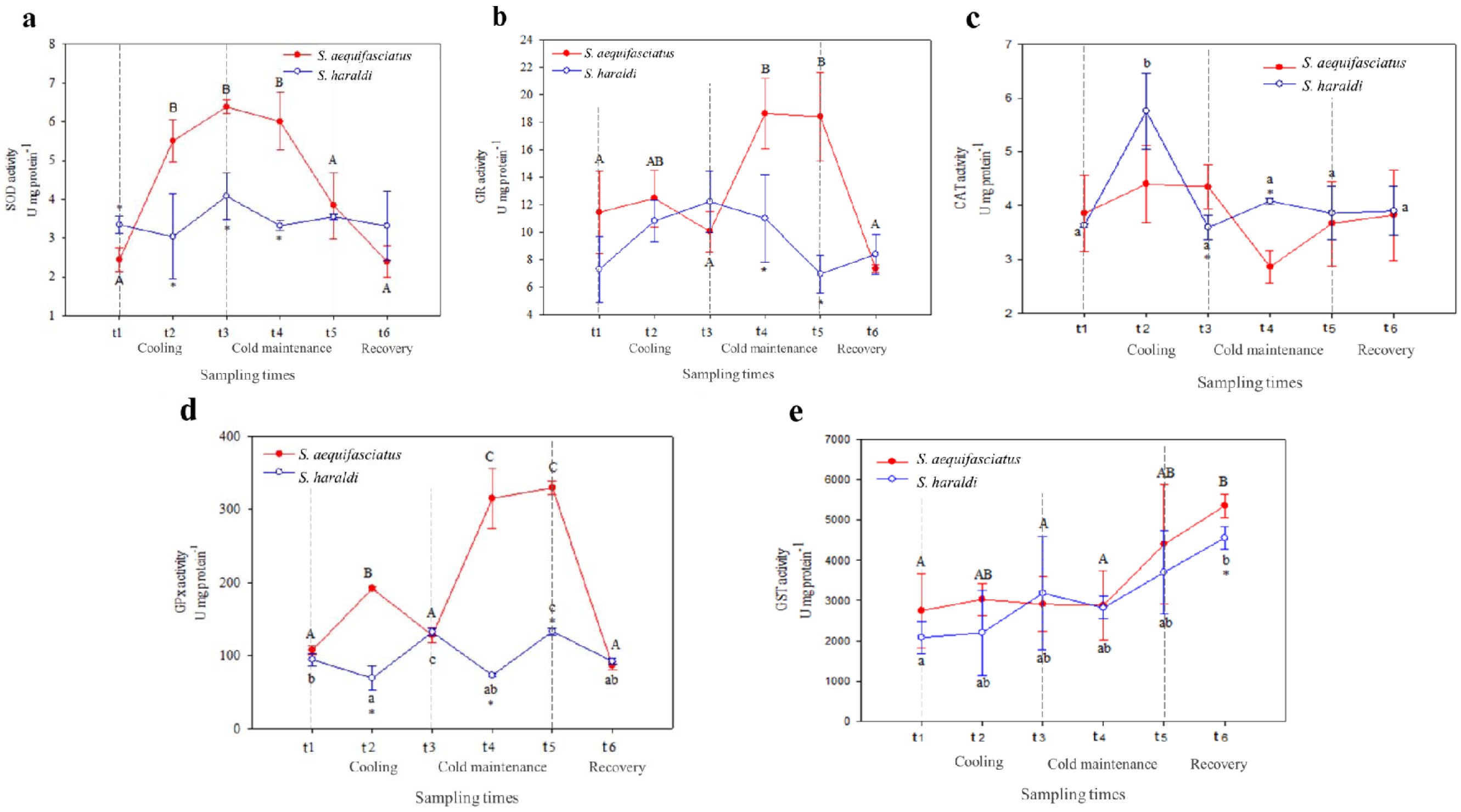
Activities of SOD (a), GR (b), CAT (c), GPx (d) and GST (e) in the gills of two subspecies of *Symphysodon spp*. were measured. red line, *S. aequifasciatus* and blue line, *S. haraldi*. Data are presented as means ±SD (n=3). *indicates significant differences (*p<*0.05) between subspecies. Different uppercase letters indicate significant differences (*p<*0.05) between sampling times within the *S. aequifasciatus*. Different lowercase letters indicate significant differences (*p<*0.05) between sampling times within the *S. haraldi*.

### 3.3 Glutathione contents

Neither *S. aequifasciatus* nor *S. haraldi* showed significant response under cold stress on GSH content (Fig. 4). However, the latter value was always significantly higher than the former value.

**Fig. 4.**
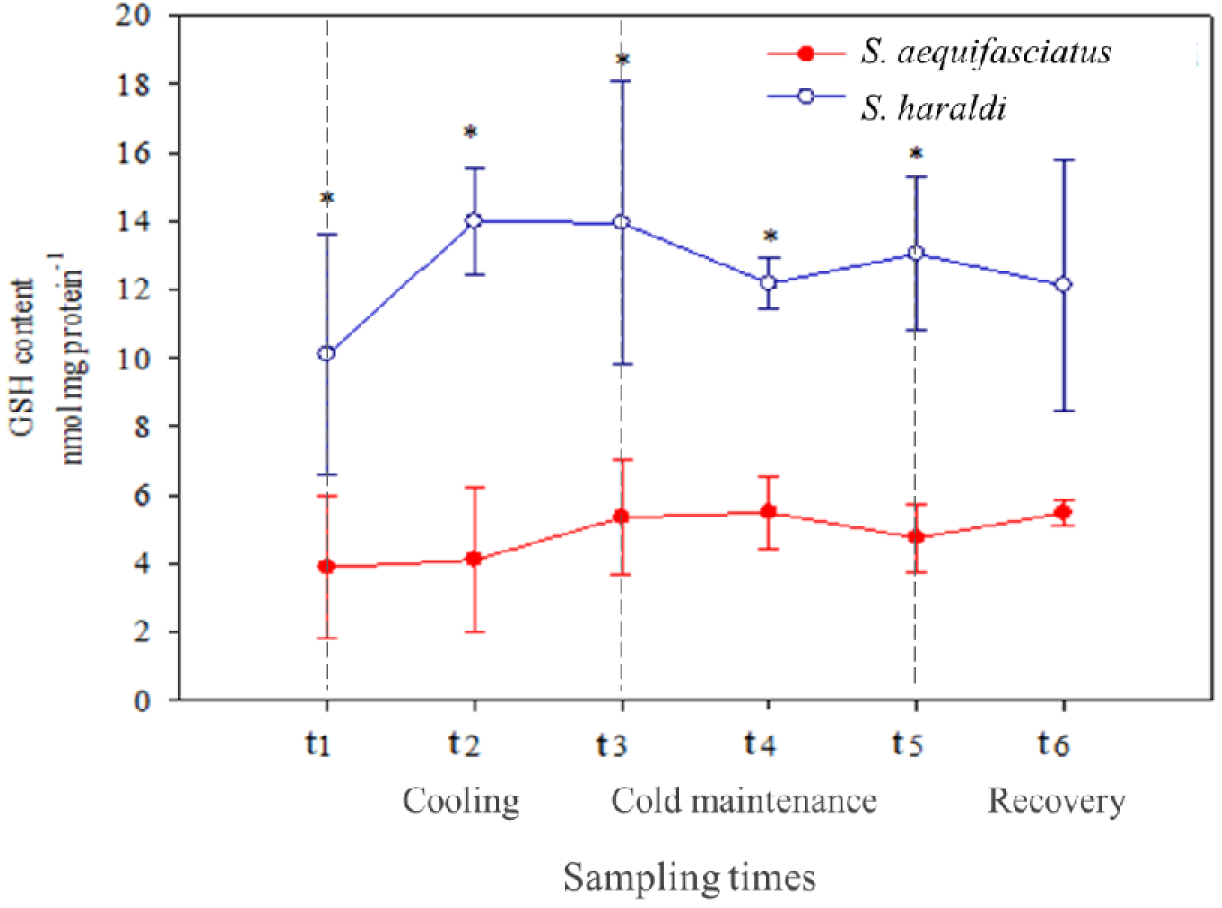
Level of reduced GSH in the gills of two subspecies of *Symphysodon spp*. red line, *S. aequifasciatus* and blue line, *S. haraldi*. Data are presented as means ±SD (n=3). *indicates significant differences (*p<*0.05) between subspecies. Different uppercase letters indicate significant differences (*p<*0.05) between sampling times within the *S. aequifasciatus*. Different lowercase letters indicate significant differences (*p<*0.05) between sampling times within the *S. haraldi*.

### 3.4 Oxidative damage

During cooling, there were no significant effects on PC (Fig. 5a) and MDA (Fig. 5b) content in both species. During cold maintenance, an increase in PC content were found in *S. aequifasciatus*, and MDA content significant increase in both species. During recovery, MDA content remained at highest level in *S. aequifasciatus*, but decreased in *S. haraldi*. PC content was not affected in both species during recovery.

**Fig. 5.**
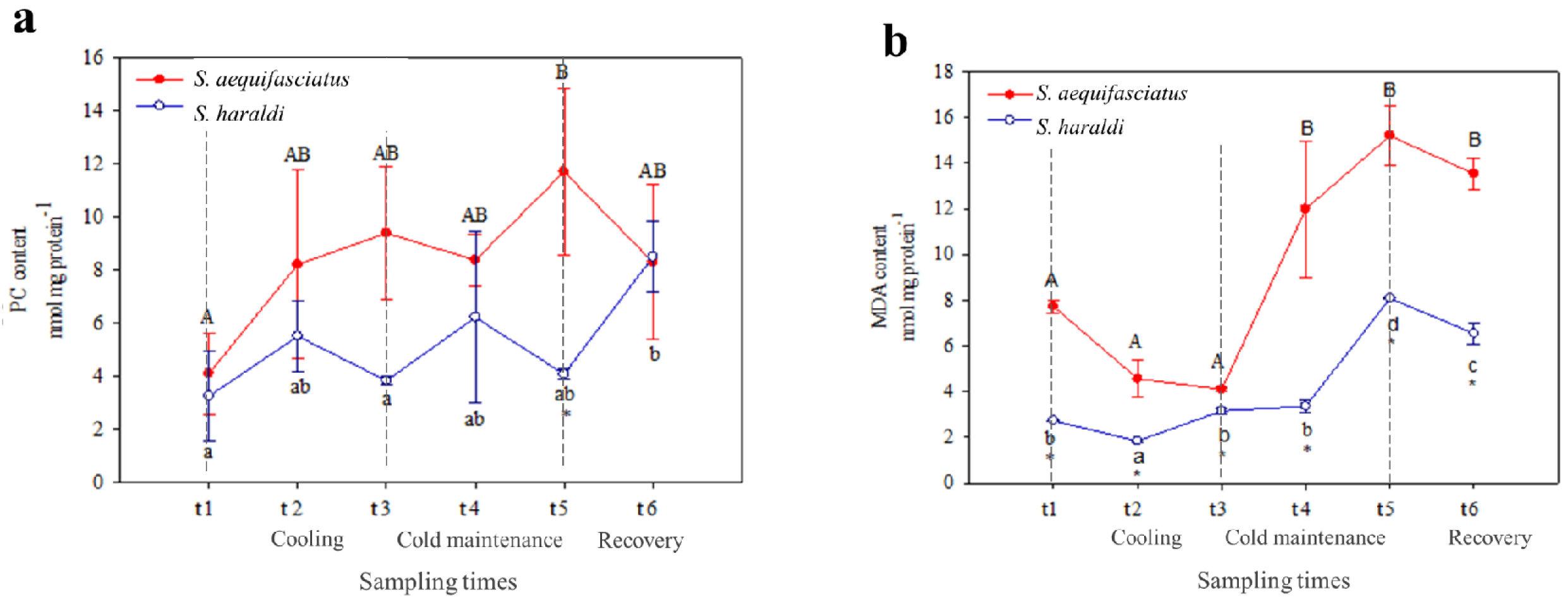
Levels of PC (a) and MDA (b) in the gills of two subspecies of *Symphysodon spp*. red line, *S. aequifasciatus* and blue line, *S. haraldi*. Data are presented as means ±SD (n=3). *indicates significant differences (*p<*0.05) between subspecies. Different uppercase letters indicate significant differences (*p<*0.05) between sampling times within the *S. aequifasciatus*. Different lowercase letters indicate significant differences (*p<*0.05) between sampling times within the *S. haraldi*.

### 3.5 Correlation analysis and comprehensive analysis on hardiness indexes for different discus fish species

Table 3 showed that the GPx, GR, GSH, GST, SOD activities had positive correlation with ROS content, and ROS content had negative correlation with CAT activity and MDA, PC content. PC content had significant and negative correlation with SOD activity. MDA content had significant and positive correlation with CAT and SOD activities, while had significant and negative correlation with GPx and GST activities.

**Table 1.** Correlation analyses among cold-resistance indices

| Index | GPx | CAT | GR | GSH | GST | SOD | MDA | PC | ROS |
| --- | --- | --- | --- | --- | --- | --- | --- | --- | --- |
| GPx | 1 |  |  |  |  |  |  |  |  |
| CAT | -0.724* | 1 |  |  |  |  |  |  |  |
| GR | 0.796* | -0.765* | 1 |  |  |  |  |  |  |
| GSH | -0.188 | 0.479* | -0.035 | 1 |  |  |  |  |  |
| GST | 0.63* | -0.855* | 0.825* | -0.079 | 1 |  |  |  |  |
| SOD | -0.032 | 0.622* | -0.087 | 0.694* | -0.495* | 1 |  |  |  |
| MDA | -0.75* | 0.709* | -0.391 | 0.652* | -0.489* | 0.524* | 1 |  |  |
| PC | -0.042 | -0.305 | -0.164 | -0.163 | 0.408 | -0.719* | -0.4 | 1 |  |
| ROS | 0.727* | -0.218 | 0.626* | 0.01 | 0.301 | 0.329 | -0.425 | -0.373 | 1 |

\* denote significant at the 0.05 probability levels

**Table 2.** The eigenvalues, proportions and cumulative of principal components

| Principal component | Eigenvalue | Proportion % | Cumulative % |
| --- | --- | --- | --- |
| Z1 | 4.493 | 49.921 | 49.921 |
| Z2 | 2.495 | 27.723 | 77.644 |
| Z3 | 1.139 | 12.658 | 90.302 |

**Table 3.**
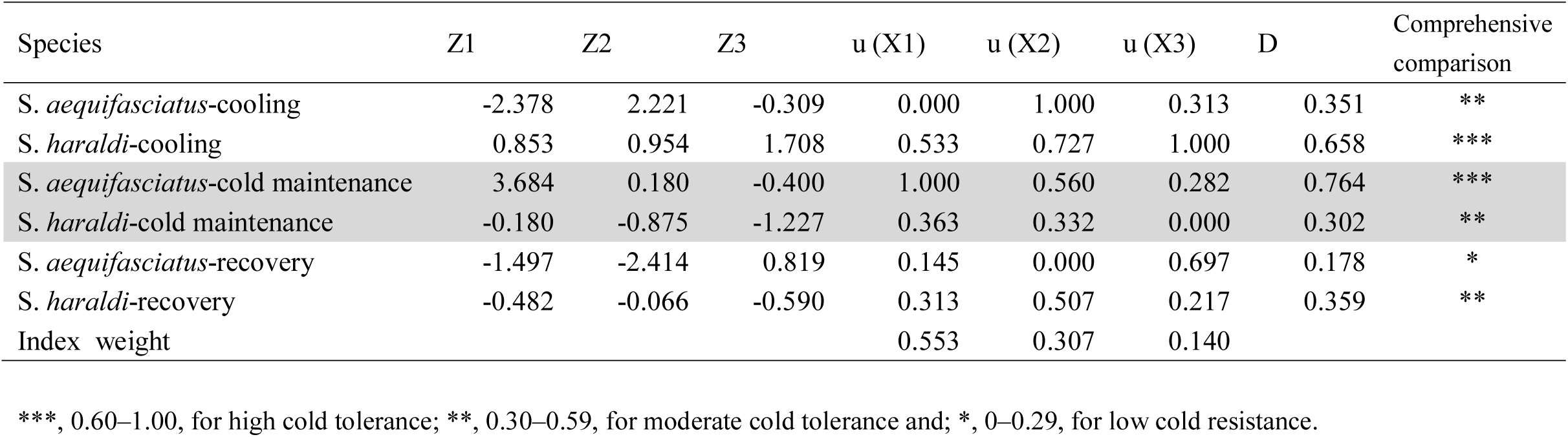
The value of comprehensive index [Zi], index weight, u (Xi), D value and comprehensive valuation for each treatment of two species.

The heat map synthesize the expression values of all hardiness indexes for the two species during cold treatment (Fig. 6). The value of hardiness indexes in discus fish differed in different species under different temperature treatment, as evident from the intensity of colors (level of expression). In *S. haraldi*, GSH activity was the only index sustained up-regulated throughout the treatment, meanwhile GST activity was significantly up-regulated at the end. But in *S. aequifasciatus*, most of the hardiness indexes were seriously affected by low temperature, and significantly up-regulated, such as SOD, GPx, GR, GST activities and ROS, PC, MDA contents.

**Fig. 6.**
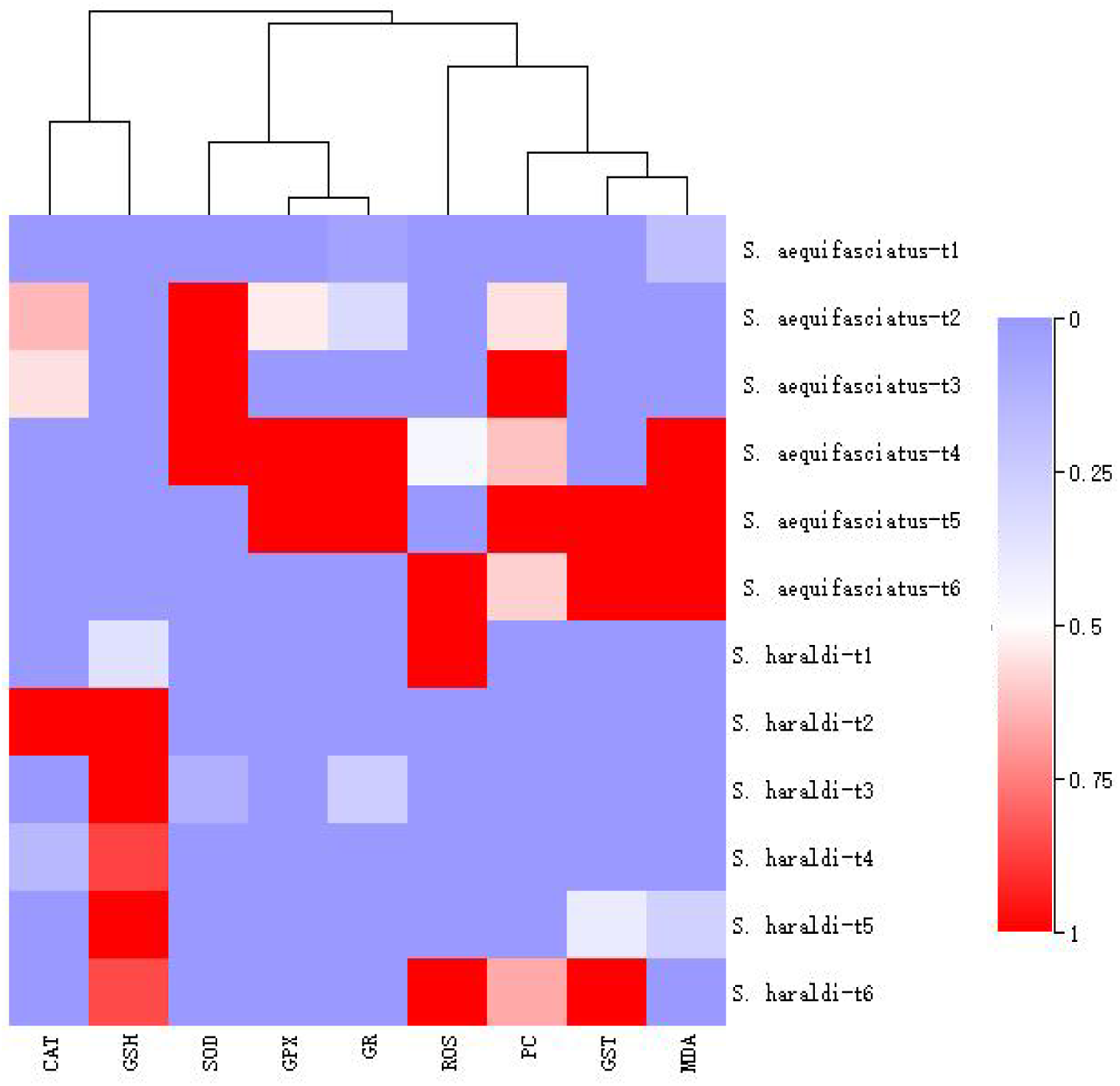
Heat-map visualization of the differential biomarkers of oxidative stress in response to cold stress between two species. Colour denotes the abundance of biomarkers of oxidative stress, from the highest (red) to the lowest (blue).

### 3.6 Using Principal component analysis (PCA) and Subordinate Function (SF) to Evaluate Hardiness of two species

As can be seen from Table 2, the first, second, and third principal component variance contribution rates reached 49.921, 27.723, and 12.658%, respectively. Notably, the cumulative variance contribution rate was 90.302% (more than 85%) without missing variables. Therefore, the first three principal components can reflect completely the different information of the cold resistance system and most of the data had already been included in the three principal components.

Subordinate function values of various comprehensive indicators of each treatment were calculated in accordance with Equation (1) (Table 3) following PCA. For the same comprehensive indicator, such as Z1, the maximum u (X1) was 1.000 for the *S. aequifasciatus*-cold maintenance treatment and 0.000 for *S. aequifasciatus*-cooling. This suggested that when only Z1 was considered, *S. aequifasciatus* showed the highest level of adaptability to the cold maintenance treatment, whereas its adaptability to the cooling treatment was the lowest. The adaptabilities to the remaining treatments were sorted according to the value of u (Xi).

Based on the contribution of various comprehensive indicators, the weights were calculated in accordance with Equation (2). The results showed that the three comprehensive indicators had weights of 0.553, 0.307 and 0.140, respectively (Table 3).

The comprehensive physiological adaptability capabilities of two species to various water temperature were calculated in accordance with Equation (3) (Table 3), and sorted based on the value of D. The higher D value, the higher cold hardiness showed. To be specific, the cold hardiness of *S. haraldi* during cold treatment was in the order of cooling > cold maintenance ≈ recovery, and the cold hardiness of *S. aequifasciatus* during cold treatment was in the order of cold maintenance > cooling > recovery. It is important to note that the minimum D value was obtained for *S*. *aequifasciatus* during recovery, suggesting the lowest cold hardiness, whereas the cold hardiness based of *S. aequifasciatus* during cold maintenance and for *S. haraldi* during cooling were classified as highest level.

## 4. Discussion

### 4.1 Oxidative stress levels increased in both species by acute cold stress

Many studies have found that an acute temperature decrease has an influence on haematological and metabolic processes (Ban, 2000; Sun et al., 1995), which could promote the generation of ROS (Joe 2017; Martínez-Álvarez et al., 2005). This is agree with present finding in two discus fish. Under oxidative stress, antioxidant defense system inhibiting an excess of oxyradical formation (Joe., 2017; Ren et al., 2018; Vinagre et al., 2012; Ye et al., 2015). For example, Malek (2004) found that SOD and GPx isoforms and thioredoxin, but not CAT, upregulated in zebrafish skeletal muscle under cold stress. Similarly, our previous studies also found that when *S. aequifasciatus* under chronic cold stress, the activities of SOD and GPx, and level of GSH increased while the production of ROS increased, but the production of MDA not increased (Wen et al., 2018). Unlike chronic cold stress, acute cold stress caused oxidative damage on both species, revealed by the increased level of MDA and PC (Almroth et al., 2008; Enzor and Place, 2014; Trenzado et al., 2006). Due to the antioxidative index was directly correlated to temperature, it therefore appears that oxidative stress levels could provide information on cold hardiness of fish.

### 4.2 Is oxidative stress higher in *S. aequifasciatus* than *S. haraldi*?

The excess ROS leading to oxidative stress in fish (Atli et al., 2016; Joe, 2017; Martínez-Álvarez et al., 2005). Therefore, the present study showed that the ADS might be able to successfully prevent oxidative stress in *S. haraldi*, but in *S. aequifasciatus* (Ates et al., 2008; Atli and Canli, 2007; Eyckmans et al., 2011).

The ADS, such as SOD, CAT, GPX and GR, usually act in a coordinated manner in order to ensure the optimal protection against oxidative stress (Morales-Medina et al., 2017). Following temperature reduction, SOD and GPx activities also upregulated in zebrafish skeletal (Malek et al., 2004). The research in cunner (*Tautogolabrus adspersus*) also found that fish acclimated to cold temperature had higher levels of GR transcript in both the head kidney and liver (Alzaid et al., 2015). Attributed to complementary activity of GPX to CAT activity, CAT activity usually showed increased trend while a decreasing trend was observed for GPX (Atli and Canli, 2010; Saglam et al., 2014; Santovito et al., 2012). GSH also can neutralise ROS, playing an important role as a cofactor for various glutathione-dependent antioxidant enzymes (Grim et al., 2013; Halliwell and Gutteridge, 2007; Sedlak and Lindsay, 1968). For example, Heise et al. (2007) found that GSH content was two to three times more concentrated in polar compared to temperate eelpout liver, suggesting that polar eelpout are more susceptible than their North Sea confamilials. Klein et al (2017) putted forward an idea that the higher SOD and CAT activity observed in peripheral tissues of *N. rossii* respect with *N. coriiceps* might showed the former need a more powerful ADS than the latter fish species. In this case, it seems that *S. aequifasciatus* was more susceptible than *S. haraldi*, and needs more powerful ADS.

But at the same time, MDA content, as oxidative damage marker (Joy et al., 2017; Ren et al., 2018; Vinagre et al., 2012; Ye et al., 2016) was significantly higher in *S. aequifasciatus* than in *S. haraldi*. It suggested that *S. haraldi* better protected from oxidative damage than *S. aequifasciatus*.

From the above, oxidative stress higher in *S. aequifasciatus* than *S. haraldi* under acute cold stress.

### 4.3 The reason of species-specific cold resistance between *S. aequifasciatus* and *S. haraldi*

In addition to their Amazon basin-wide distribution, different environmental pressure were subjected by discus fish in different geographic gradients (Eliason et al., 2011; Heise et al., 2007; Troia and Gido, 2016 and 2017;). Noteworthily, *S. aequifasciatus* distributes at the upstream of Amazion River, and *S. haraldi* distributes at the midstream and downstream, while both species distribute at the similar latitudinal gradients (Ready et al., 2006). A recent research by Carmona-Catot (2011) found that upstream-to-downstream gradients are as influential as latitudinal gradients in shaping growth, reproduction, and body condition among European populations of *Gambusia holbrooki*. And Model results from the Madison River in Montana indicate that, on average, rainbow trout at the downstream site (B) would have a stress index that is 2±3 times greater than rainbow trout at the upstream site (A) even though the difference in mean temperature is only 0.48 °C (Bevelhimer and Bennett, 2000). It suggested that *S. haraldi* had a greater stress index than *S. aequifasciatus* likely contribute to the upstream-to-downstream gradients.

Environmental pressure has led fish in the region to develop considerable genomic plasticity during their evolutionary process, and a series of ecological, morphological, physiological, metabolic and molecular adjustments can be seen. Indeed, the analysis of mitochondrial DNA haplotypes, chromosomal complement and meiotic organization indicates that the western Amazonian *Symphysodon*, *S. aequifasciatus*, showed interspecific variability from the central Amazonian *Symphysodon*, *S. haraldi* (Gross et al., 2009; Gross et al., 2010; Gross et al., 2006; Ready et al., 2006). These adjustments might help them to maintain organic homeostasis and allow them to survive during these environmental changes. According to Chippari-Gomes (2003), *Symphysodon* species positively exhibited different adapt capacity, which allows them to survive in conditions of moderate hypoxia. Place et al. (2004) found that the loss of the HSR in the Antarctic notothenioids resulted in the inability of *T. bernacchii* to upregulate hsp70 mRNA during a 1 h in vitro thermal stress at temperatures as high as +10°C. In contrast to the loss of the HSR in the notothenioids, *Lycodichthys dearborni*, a phylogenetically distant Antarctic species, has retained the ability to upregulate the expression of the hsp70 gene in response to thermal stress (Place and Hofmann 2005). In view of these, there should be a more extensive study using molecular methodologies to clarify the genetic variability which correlated with adaptation to temperature among two species.

## 5. Conclusion

The ROS generation, ADS and oxidative damage can be used as hardiness indexes in *Symphysodon*. *S. haraldi* which is found in the central portion of the Amazon basin show a stronger cold resistance than *S. aequifasciatus* which is found in the west portion of the Amazon basin, exhibiting a significant interspecific variability under acute cold stress.

## Acknowledgements

The authors are grateful to anonymous referees for professional judgment of the manuscript that helped substantially improve the work.

## Competing interests

We have no competing interests.

## Author contributions

B.W. drafted the paper, S.R.J. conducted the measurement and analysis, Z.Z.C. and J.Z.G. designed the research, L.W. conducted the Methodology, H.P.L. conducted the animal culture, Y.L. conducted the sampling procedures.

## Funding

The study presented in the manuscript was funded by the Shanghai Sailing Plan for the Young Scientific Talents (19YF1419400).

## Supplementary information

This article has no supplementary information.

